# Optimization of the prostaglandin F_2α_ receptor for structural biology

**DOI:** 10.1101/2025.02.15.638479

**Authors:** Marine Salze, Sébastien Chrétien, Tegvir S. Boora, Madalina Macovei, Eric Barbeau, Véronique Blais, Stéphane A. Laporte, Martin Audet

## Abstract

Prostaglandin F_2ɑ_ (PGF_2ɑ_) is a bioactive lipid derived from arachidonic acid and is involved in many physiological and pathophysiological processes such as parturition, vascular tone regulation, glaucoma and inflammation. It acts by binding to the Prostaglandin F_2ɑ_ receptor (FP), a G Protein-Coupled Receptor (GPCR) that mediates signaling events by engaging intracellular heterotrimeric G protein effectors. The orthosteric binding site of lipid-binding receptors displays greater efficacy-dependent plasticity that hinders the design of ligands. Solving the structure of FP with ligands of different efficacies at an atomic level is important to fully understand its mechanism of activation and inhibition. Most purified FP-ligand complexes are unstable *in vitro*. The development of new X-ray crystallography and single particle cryo-electron (cryoEM) strategies to understand receptors’ signal transduction requires improved purification yield and *in vitro* stability of the receptor. Here, we present a protein engineering effort to optimize the FP protein sequence for use in structural biology. Strategies involve protein insertion sites in the third intracellular loop (ICL3), N-terminal and C-terminal deletions, and single-point mutations that favorably affect receptor purification yield and stability *in vitro*. The best FP construct displays a yield of 1.5 mg/L and a stability of 59°C that constitute a threefold improvement in purification yield and 9°C increase in stability over the wild-type receptor. These modifications in the receptor are suitable for pursuing alternative strategies for improving FP purification yield and for studying FP binding efficacy to its ligands through structural biology approaches.

## Introduction

G Protein-Coupled Receptors (GPCRs) form the largest family of membrane proteins in the human genome and are the target of more than 35 % of drugs in Western countries [1, 2]. Prostaglandin F_2α_ (PGF_2α_) is a bioactive lipid that binds and activates FP, a GPCR at the cellular plasma membrane, leading to the engagement of intracellular heterotrimeric G proteins effectors and subsequent activation of cellular signaling cascades [3, 4]. PGF_2α_-mediated activation of FP plays an important role in many physiological processes and pathological conditions such as parturition, vascular tone regulation, intracerebral hemorrhage, atherosclerosis, ocular hypertension and glaucoma [4–9], making it an important pharmaceutical target. As such, PGF_2α_ derivatives are currently used in clinics. For example, FP activator (agonists) such as fluprostenol and latanoprost are used for the treatment of glaucoma and ocular hypertension [4]. Cloprostenol, the most potent FP ligand, is used for labor induction [4]. Few FP inhibitors (antagonists) have been developed, such as AL8810 [10], PDC113.824 (a positive allosteric modulator biased toward the Gq pathway) [11], BAY-6672 [9], and OBE022, which showed encouraging outcomes in clinical trials for addressing preterm labor [12]. Overall, the development of antagonists targeting FP remains a challenge [13].

Recently, the development of new approaches in structural biology has led to high-resolution structures of FP-bound PGF_2α_ and closely related ligands, providing the first insight into agonist binding to the receptor [14, 15]. In these structures, FP shares the typical GPCR topology consisting of 7 transmembrane helical domains, 3 intracellular loops, 3 extracellular loops, a helix 8 parallel to the membrane, and extended N- and C-terminal tails. The receptor folds as a 7 transmembrane helical bundle with PGF_2α_ binding at the orthosteric site, inside the bundle, covered by an extended N-terminus that serves as a lid to the receptor. This feature, shared by many lipid binding receptors, contributes to the typical low off-rate of lipid receptor ligands and in the difficulty to determine their pharmacological profiles [14–17]. In addition, the greater efficacy-dependent plasticity of lipid receptor binding site complicates the rational design of antagonists [16]. A full understanding of the mechanisms of activation and inhibition of FP requires solving structures with ligands of different efficacies.

Stabilization and improvement of the purification yield of GPCRs through optimization of receptor sequences have been key features in the recent development of GPCR structural biology derived from cryo-electron microscopy (cryoEM) and X-ray crystallography approaches [18–20]. So far, FP could only be stabilized by the binding of an agonist and in complex with a chimeric heterotrimeric G protein and stabilizing nanobody [14, 15]. In addition, FP has never been crystallized, and apo or antagonist-bound receptor are unstable *in vitro.* This difficulty is shared among some of the prostaglandin receptor, as there is only one co-crystal structure with an antagonist, the PGE_2_ type 4 receptor bound to ONO-AE3-208 [21]. This structure required extensive receptor optimization and the use of a Fab antibody to stabilize the antagonist-bound conformation of the receptor. To foster further structural studies of FP and gain a full understanding of the ligand’s mechanism of efficacy, it is imperative to improve methods to overcome its lack of stability during purification. Here, we present work on the optimization of the receptor protein sequence to improve the purification yield and stability of the FP receptor *in vitro*.

## Materials and Methods

### Cloning and gene modification

Human FP gene sequence was codon optimized for *Sf9* and human cells expression, synthesized by GeneScript (Piscataway, USA) and cloned into the pFastBac® vector with a FLAG tag, 10xHis tag and TEV cleavage site at the N-terminus. Insertion of soluble domains, receptor truncation and single residue mutation were performed by PCR using Phusion® polymerase (ThermoFisher Scientific, Waltham, USA). For expression in HEK293T cells, the tagged receptor constructs were subcloned into pcDNA3.1(-) using BamHI-HF/HindIII-HF and T4DNA ligase (New England Biolabs).

### Expression and purification of FP

FP constructs were expressed in *Sf9* insect cells using the Bac-To-Bac Baculovirus expression system (ThermoFisher Scientific). *Sf9* insect cells line is a clonal isolate derived from the parental *Spodoptera frugiperda* cell line IPLB-Sf-21-AE purchased directly from Expression Systems (cat#94-001F). They were grown in ESF 921 Insect cell culture protein free media (Expression Systems, Davis, USA) at 27°C with shaking. Membranes from cells expressing FP constructs were prepared using two rounds of washing and centrifugation at 45,000 X g, first in the presence of lysis buffer containing 10 mM HEPES, pH 7.5, 20 mM KCl, and 10 mM MgCl_2_, and then with a washing buffer containing 10 mM HEPES pH 7.5, 1 M NaCl, 20 mM KCl and 10 mM MgCl_2_. Protease inhibitors (1 mM Benzamidine and 1 µg/mL leupeptin) were added in both steps. The purified membrane was then resuspended in a storage buffer (10 mM HEPES pH 7.5, 20 mM KCl, 10 mM MgCl_2_, 20% (v/v) Glycerol). Membrane containing receptor was then solubilized in 50 mM HEPES pH 7.5, 800 mM NaCl, 5 mM KCl, 2.5 mM MgCl_2_, 0.5% (w/v) n-dodecyl-β-D-maltopyranoside (DDM, Anatrace), and 0.1% (w/v) cholesteryl hemisuccinate (CHS, Sigma) in the presence of protease inhibitors, 1 mg/mL iodoacetamide and 10 µM cloprostenol (Cayman chemical, USA). The supernatant was isolated by centrifugation for 1 h at 21,000 X g and then incubated with TALON resin (TALON® Superflow™ beads, Cytiva) overnight at 4°C. The TALON resin was washed with 20 column volumes of wash buffer 1 containing 50 mM HEPES pH 7.5, 150 mM NaCl, 20 mM MgCl_2_, 20 mM imidazole, 8 mM ATP, 10% (v/v) glycerol, 0.1% (w/v) DDM, 0.02% (w/v) CHS, and 1 µM cloprostenol, followed by 10 column volume of wash buffer 2 containing 50 mM HEPES pH 7.5, 150 mM NaCl, 30 mM imidazole, 10% (v/v) glycerol, 0.05% (w/v) DDM, 0.01% (w/v) CHS, and 1 µM cloprostenol. The receptor was eluted using 6 column volumes of elution buffer containing 50 mM HEPES pH 7.5, 150 mM NaCl, 250 mM imidazole, 10% (v/v) glycerol, 0.015% (w/v) DDM, 0.003% (w/v) CHS, and 1 µM cloprostenol. Finally, the purified receptor eluate was concentrated using a vivaspin 500, 100 Kd (Sartorius), and analyzed by SEC on an Infinity II HPLC system, (Agilent) using an SRT-C SEC-300 column (Sepax). Purification yield was determined using UV_280_ absorbance peak at the elution time corresponding to the monodispersed state of the receptor on HPLC-SEC chromatogram.

### Microscale fluorescence thermal assay

CPM dye was obtained from ThermoScientific and resuspended in DMSO at a concentration of 500 µM. 2 µg of purified protein was added to the reaction buffer (50 mM HEPES pH 7.5, 150 mM NaCl, 0.05% (w/v) DDM, 0.01% (w/v) CHS, and 5 µM CPM). The samples were then treated with 100 µM cloprostenol or vehicle and incubated for 20 min on ice. Unfolding of FP was induced using a Rotor-Gene Q (QIAGEN) thermocycler by slowly increasing the sample temperature (+0.5°C / step from 30°C to 95°C), resulting in an emission of a fluorescence signal detected in the blue channel (excitation 365nm/emission 460nm) from the CPM chemical reaction to newly exposed receptor cysteines. The melting temperature (Tm) was determined as the temperature at the maximal value of the first derivative of the fluorescence vs temperature thermodenaturation curve.

### Cell signaling assay

HEK293T cells were obtained through ATCC (cat#CRL-3216) and cultured in DMEM supplemented with 10% FBS and 20 μg/mL gentamicin. Cells were grown at 37°C in 5% CO_2_ and 90% humidity. Cells were seeded at a density of 9,000 cells per well in a white 96 well flat bottom plate and then transiently transfected with receptor and sensor DNA 24h later. The previously described RhoA BRET biosensor was used to assess G protein response [22, 23]. All assays were performed 48 h post-transfection. On the day of the experiment, cells were incubated for 1 h with Tyrode’s buffer (25 mM HEPES, pH 7.4, 140 mM NaCl, 12 mM NaHCO_3_, 5.6 mM D-glucose, 2.7 mM KCl, 1 mM CaCl_2_, 0.5 mM MgCl_2_, 0.37 mM NaH_2_PO_4_,). Cells were stimulated with serially diluted PGF_2α_ or cloprostenol from 10^−10.5^ M to 10^−5^ M, and the signal was recorded using the Biotek Synergy 2 plate reader with filter set 410/80 nm (donor) and 515/30nm (acceptor). Cell-permeable substrate coelenterazine 400a (final concentration of 2.5 μM) was added 3.5 min prior to BRET measurements. BRET ratios were calculated by dividing the intensity of signal emitted by the acceptor over the signal of light emitted by the donor. The data was fitted to 12-point concentration response curves and analyzed for its activity.

### Data presentation and statistical analysis

Graphs were created using Graphpad Prism software. Cartoons in Fig 1 were derived from the bioicons database. Snake-plots were generated by the GPCR database (GPCRdb). Figures were mounted using Inkscape software. Statistical analyses were performed using Graphpad Prism software. Statistical significance was determined by a Student’s t test or a one-way ANOVA followed by a Dunnett post-hoc analysis.

**Fig 1.**
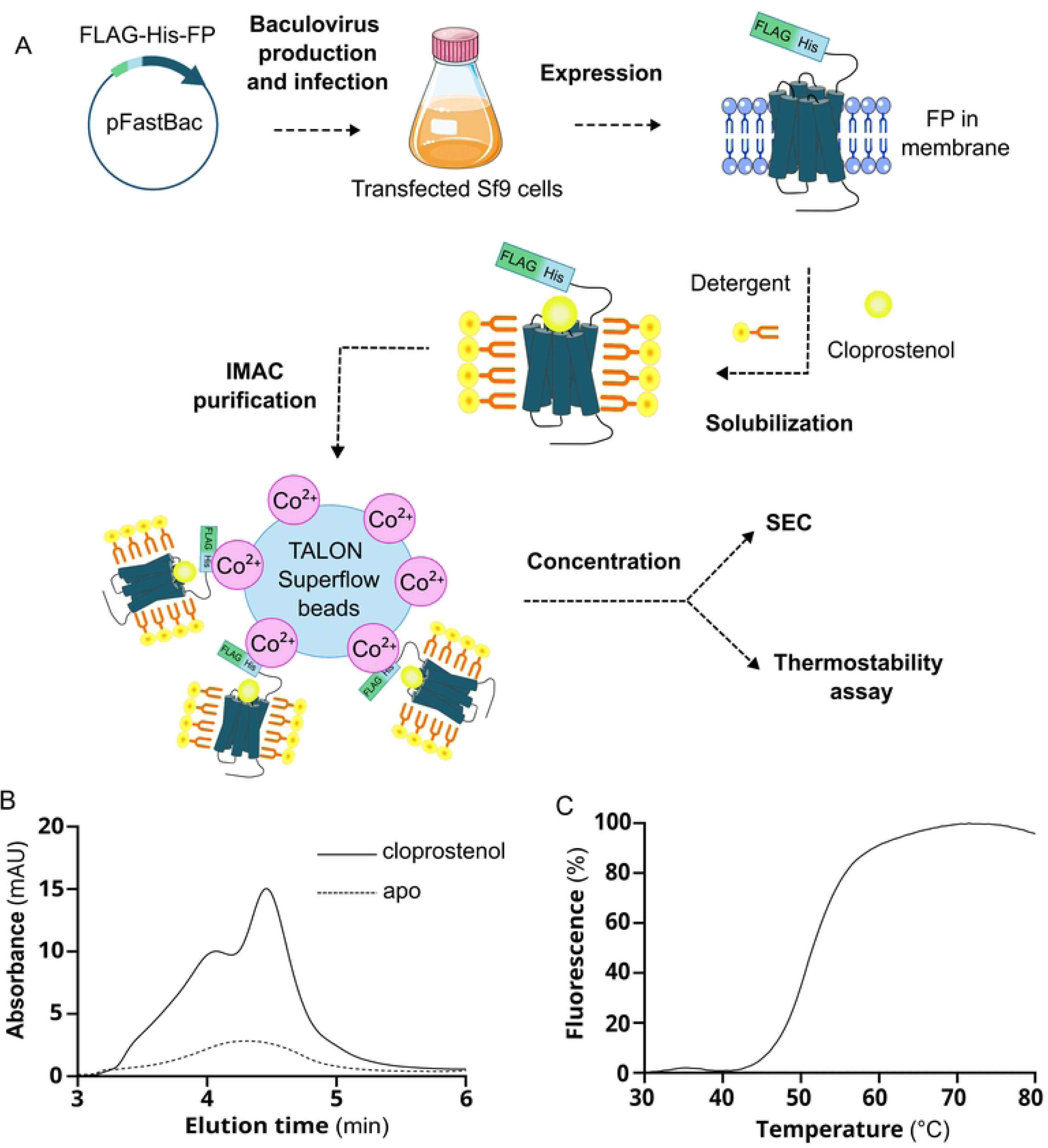
Purification of FP in complex with cloprostenol. (A) Cartoon of the steps leading to FP purification by IMAC and characterization using HPLC-SEC and microscale fluorescence thermal assay. (B) Representative HPLC-SEC chromatogram of purified FP-WT (Wild Type) in the presence and absence of cloprostenol. The mean ± SEM of FP purification yield determined from 3 independent experiments is 0.097 ± 0.008 and 0.247 ± 0.024 mg / L for the apo and the cloprostenol-bound forms, respectively. (C) Representative thermostability curve of FP-cloprostenol complex. The mean ± SEM of the Tm value determined from 3 independent experiments is 50.8 ± 0.4°C.

## Results

### FP Purification and Stability

To assess the stability of FP in micelles, we first inserted the coding sequence of the receptor fused with a FLAG and a polyhistidine (10xHis) affinity tags at the N-terminus into the pFastBac1 vector. This vector is used to produce baculoviruses that express the FP coding sequence in *Sf*9 insect cells (Fig 1A). We then purified FP from a membrane preparation of these cells using immobilized metal affinity chromatography (IMAC) and analyzed the purification yield and aggregation state of the receptor in DDM/CHS micelles using size exclusion chromatography by HPLC (HPLC-SEC) (Fig 1B). The apo form of FP displayed a low purification yield of 0.097 mg/L and a wide peak on the chromatogram with an elution time of 4.07 minutes, corresponding to a higher order aggregation state of the receptor. Since agonist-induced receptor stabilization *in vitro* has already been observed for another prostaglandin receptor and other GPCRs [17, 24], we added a saturating amount of cloprostenol, a stable PGF_2α_ analog, at all steps of FP purification. This resulted in the appearance of a major elution peak at 4.48 minutes, corresponding to the monodispersed form of the receptor. Overall, the monodispersed form of the FP-cloprostenol complex showed a 2.5-fold increase in the purification yield compared to the FP apo aggregated form (0.247 mg/L vs 0.097 mg/L, respectively). We further measured the cloprostenol-induced stability of the receptor using a microscale thermal stability assay on the FP-cloprostenol purified complex (Fig 1C). Upon heat-induced denaturation, proteins expose embedded cysteine residues that react with the 7-diethylamino-3-(4-maleimidophenyl)-4-methylcoumarin (CPM) dye, leading to fluorescence [25]. The inflection point of the thermodenaturation curve (fluorescence vs. heat) corresponds to the melting temperature (Tm) of the receptor, providing a direct measure its stability. FP-cloprostenol complex shows a Tm of 50°C, which is below the typical 60°C Tm required for structural biology studies and suggests the requirement of further optimization. Given the positive impact of cloprostenol on FP purification yield and stability, we included the ligand for all further purification steps described in this study. Our objective is to further enhance the stability and purification yield of the FP-cloprostenol complex through targeted FP protein sequence optimization.

### Stabilization of FP by bRIL insertion

Insertion of small soluble domains into GPCR intracellular loops has already been used to help their stabilization and to improve the purification yield [18, 20]. Thus, we inserted the soluble apocytochrome b562RIL (bRIL) from *E. coli* at the N-terminus of FP and into different positions of the predicted third intracellular loop (ICL3) [18] from residue 231 to 239 (Fig 2A), and determined the receptor purification yield and stability in DDM micelles using the microscale thermal fluorescence assay (Fig 2C-D). All FP constructs were successfully expressed in *Sf*9 cells (S1 Table) and purified to at least 95% purity as shown by a Coomassie-stained protein gel of the receptors (Fig 2B). Insertion of bRIL increased the purification yield of FP for all the constructs except the fusion between residues 231-239. The construct with the insertion between residues 232-239 showed the highest purification yield, achieving a two-fold improvement in the purification yield of 0.542 mg/L compared to 0.247 mg/L for FP without bRIL insertion. Similarly, all but one construct displayed at least a noticeable increase in stability, with three out of six being statistically significant improvements. Insertion of bRIL at position 232-239 showed the best improvement in stability with an increased Tm of 4°C (from 50 to 54), as compared to FP without insertion. N-terminal bRIL fusion to FP slightly improved the purification yield but not the stability of FP, whereas ICL3 bRIL fusion tended to improve both purification yield and stability, except for the construct 231-239 that only showed an apparent but not significant increase in Tm. Overall, the insertion of bRIL between residues 232-239 of ICL3 resulted in the best improvement in both purification yield and stability of FP in micelles.

**Fig 2.**
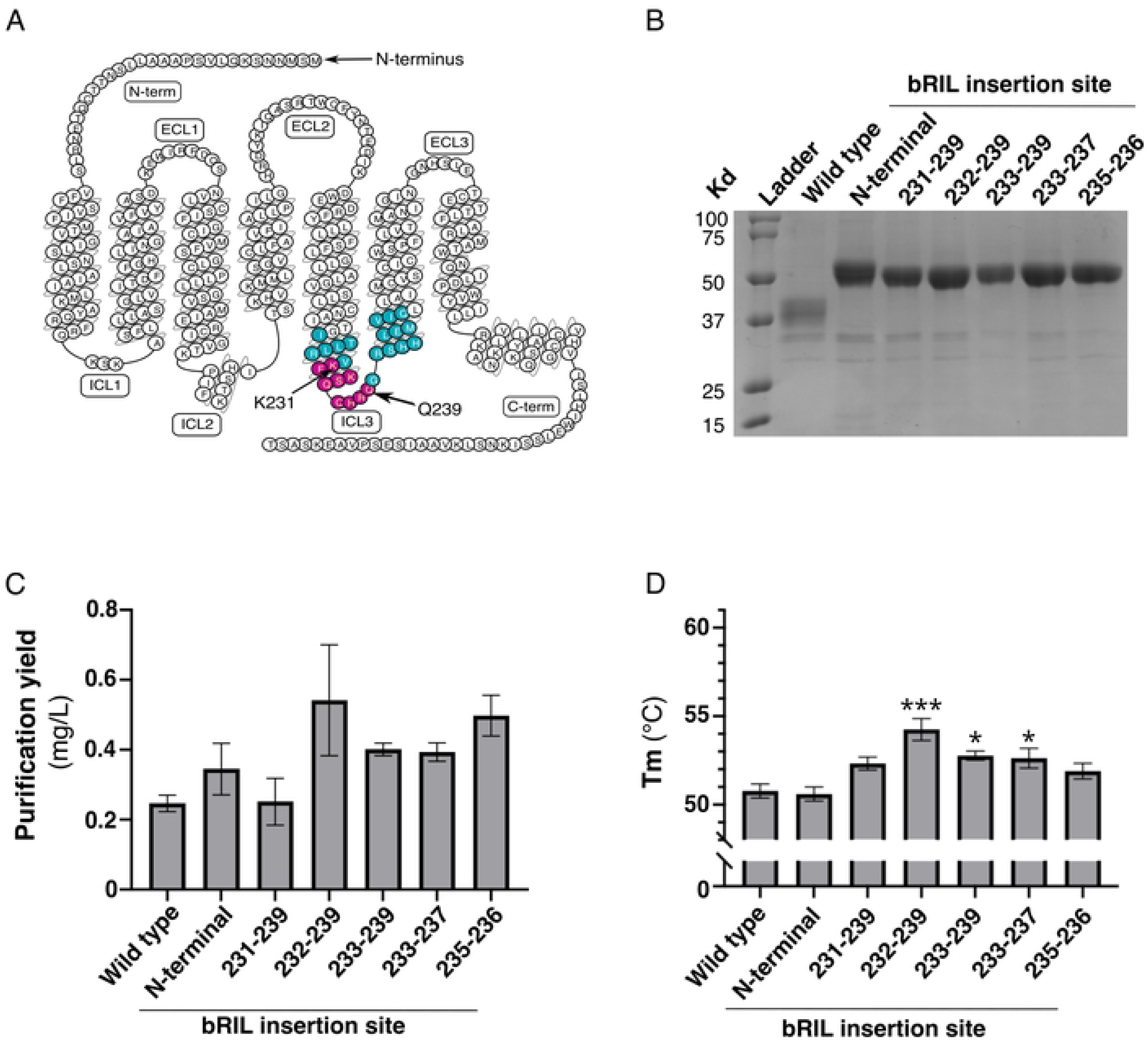
Impact of bRIL domain fusion on the stability of FP. (A) Snake plot of FP. Predicted ICL3 is indicated in blue and magenta circles. bRIL insertion in ICL3 was screened between residues 231 and 239 of FP as shown in magenta circle. (B) Coomassie-stained 12% polyacrylamide gel of the IMAC eluate of the indicated FP-bRIL fusions (C). Purification yield of bRIL fusion at the FP N-terminal, or between the residues indicated, in complex with cloprostenol. (D) Melting temperature (Tm) of the indicated FP fusions. The data are expressed as the means ± SEM of 3 independent experiments. Statistical significance was determined by one-way ANOVA followed by a Dunnett post-hoc analysis. * p-value < 0.05; *** p-value < 0.001.

### Screening of N- and C-termini deletions

We took advantage of the bRIL 232-239 insertion to further assess the impact of N-terminal truncation on FP purification yield and stability (Fig 3). The N-terminus of FP with the bRIL 232-239 insertion was incrementally deleted from residue 4 to 28, corresponding to the predicted span of the N-terminal domain in the FP cryoEM structure [14] (Fig 3A). Each construct was successfully expressed in *Sf*9 insect cells (S1 Table) and the IMAC elution fractions displayed a purity of at least 95% (Fig 3B). Deletion of fragments of the FP N-terminus generally did not affect the purification yield or the stability of FP compared to the untruncated construct (Fig 3C-D) until the truncation of residue 25, which corresponds to four residues before the beginning of transmembrane helix 1 in the FP cryoEM structure [14]. This suggests that transmembrane helix 1 may begin to form a helical turn earlier than indicated in the assigned structure, or that the few residues at the N-terminus preceding transmembrane helix 1 play a critical role in receptor folding. Overall, truncation of the N-terminus is well tolerated in *Sf*9 cells but does not provide improvement in purification yield or stability. These findings are useful in structural biology, because they allow to reduce the risk of flexible domain-induced receptor aggregation and interference with crystal contact.

**Fig 3.**
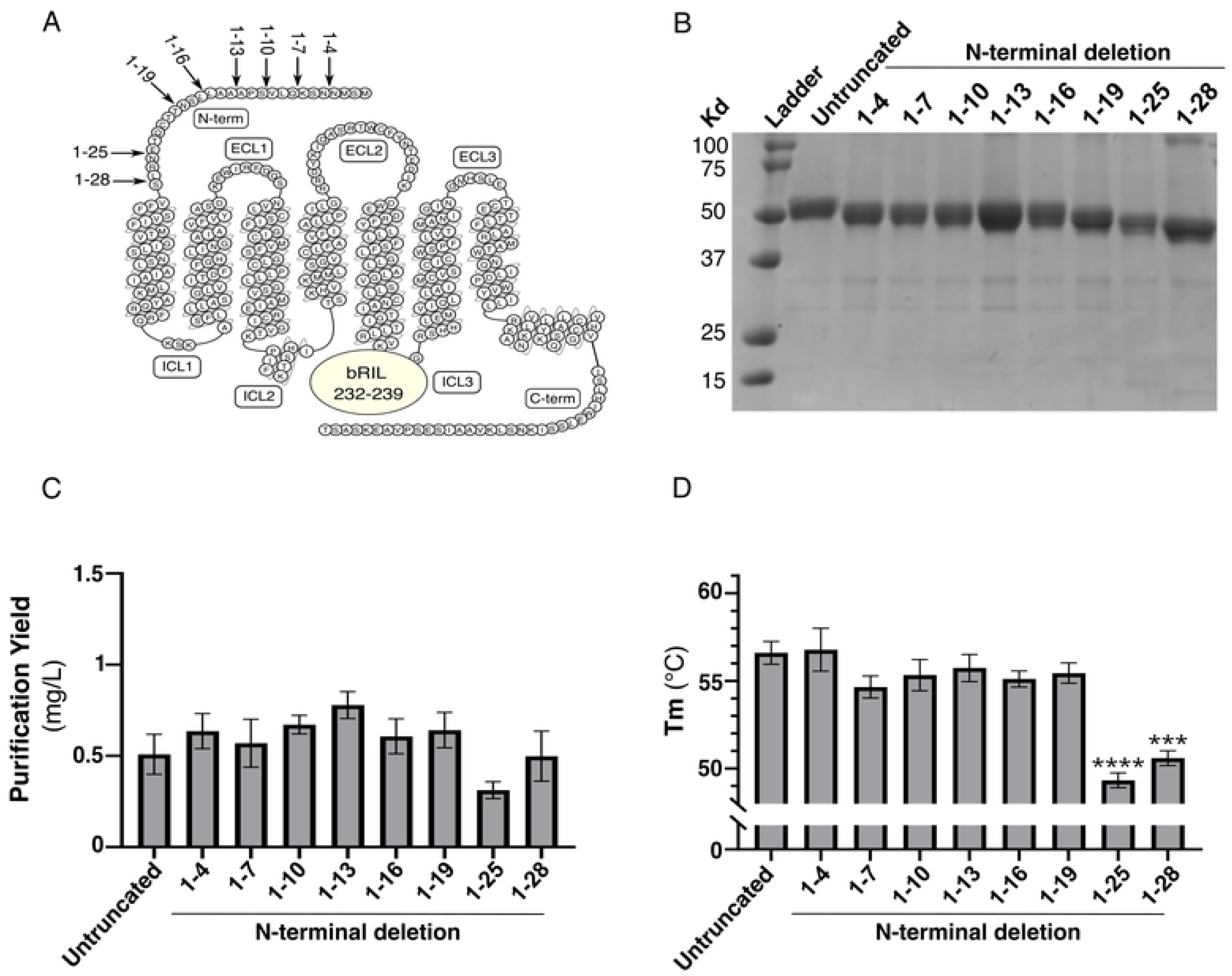
Screening of N-terminal deletion on FP stability. (A) Snake plot of FP. N-terminal deletions of the different constructs are shown by arrows. bRIL insertion in ICL3 between residues 232 and 239 is represented by a circle. (B) Coomassie-stained 12% polyacrylamide gel of the IMAC eluate of the indicated FP N-terminal deletions. (C) Purification yield and (D) melting temperature (Tm) of the IMAC eluate of indicated FP N-terminal deletions. The untruncated receptor is FP in fusion with bRIL between the receptor residues 232-239 of ICL3. The range indicates the N-terminal domain deleted from the untruncated receptor. The data are expressed as the means ± SEM of 3 independent experiments. Statistical significance was determined by one-way ANOVA followed by a Dunnett post-hoc analysis. *** p-value < 0.001; **** p-value < 0.0001.

In parallel, we investigate the effect of the C-terminal truncation on receptor purification yield and stability in the context of the N-terminal fusion of bRIL with FP (Fig 4). To achieve this, we truncated the C-terminus of the receptor every four residues up to residues 308, which corresponds to the beginning of helix 8 in the FP cryoEM structure [14] (Fig 4A). All truncated constructs expressed well in *Sf*9 cells (S1 Table). We could purify all the FP constructs up to the truncation at residue 324 (Fig 4B-C). Further truncations to residues 320, 316, 312 and 308, although well expressed in cells, could not be purified. Overall, our findings imply that truncation of the C-terminal domain does not improve the purification yield and does not affect FP stability, at least for the purified truncations up to residues 324.

**Fig 4.**
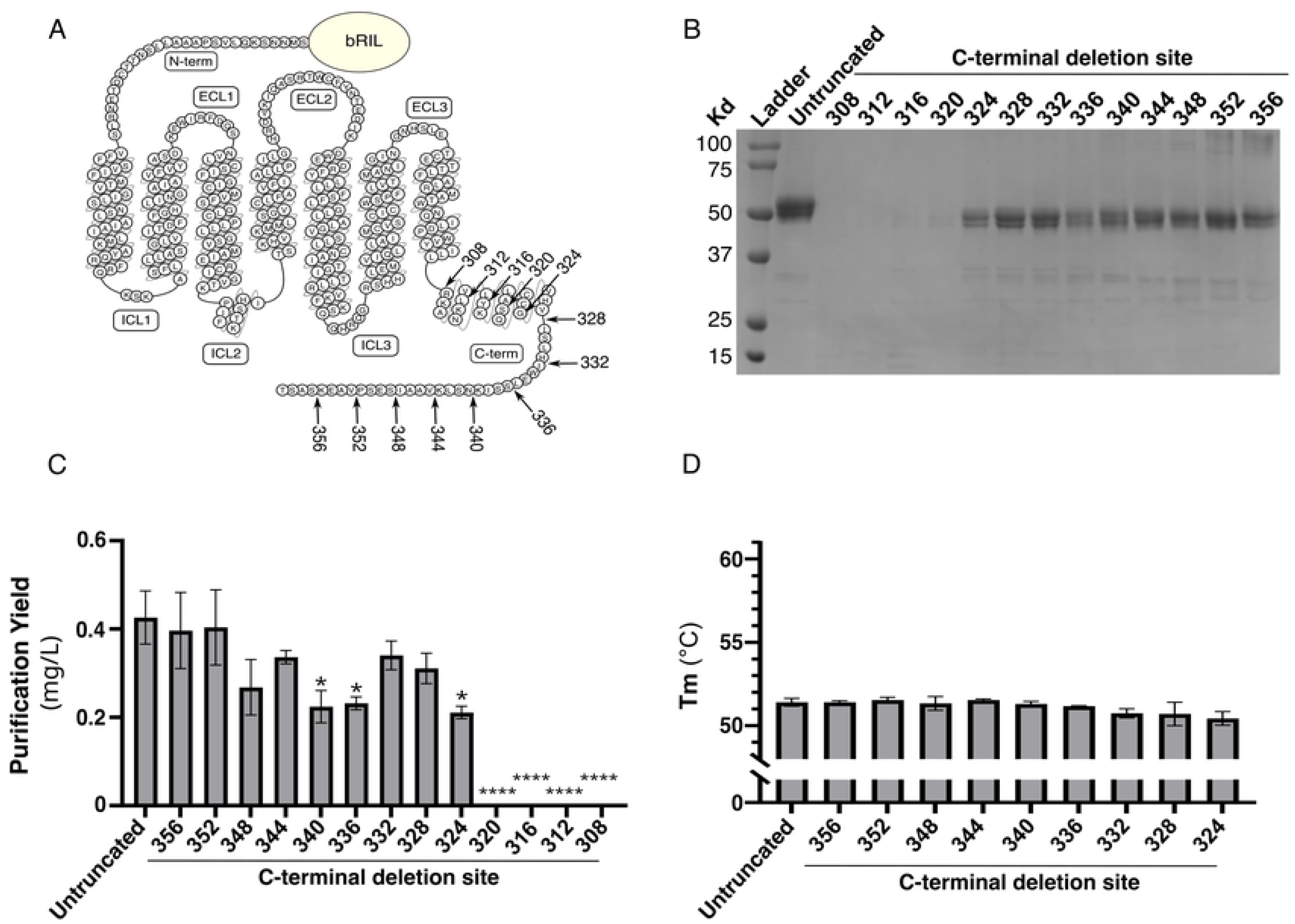
Screening of C-terminal deletion on FP purification yield. (A) Snake plot of FP. C-terminal deletion on the different constructs is shown by arrows. bRIL insertion at the N-terminus is represented by a circle. (B) Coomassie-stained 12% polyacrylamide gel of the IMAC eluates, (C) Purification yield and (D) melting temperature (Tm) of the indicated receptor C-terminal deletions. The untruncated receptor is an N-terminal fusion of bRIL with FP. The numbers indicate the last residue truncated from the C-terminus of the receptor. The data are expressed as the means ± SEM of 3 independent experiments. **** p-value < 0.0001.

### Insertion of mutations

Insertion of stabilizing mutations has been key to improving GPCR recovery during purification [26]. We thus further assessed the impact of single residue mutations on receptor stability and purification yield. We tested three mutations, A15G, S127A, and M255V, inserted on a bRIL fusion at position 232-239 of FP (Fig 5A). A15G was selected as we previously found that it increased receptor cell expression in human embryonic kidney cells. Alanine substitution at residue 127 aimed to reduce hydrophilic residues content in the membrane facing pocket formed by transmembrane helices 3-4-5, and is a similar previously used strategy that further stabilizes the crystal structure of the ß_2_-adrenergic receptor in complex with an inverse agonist [27]. Finally, M255V was selected, because it is a stabilizing mutation predicted by GPCRdb [26]. As a control, we inserted a valine at position 163 of FP, a position homologous to the V185 in the PGE_2_ type 3 receptor (EP3). In the EP3-PGE_2_ co-crystal structure, the valine at position 185 was mutated to a serine to stabilize the receptor [28]. Thus, alanine substitution of a valine at position 163 of FP would not be predicted to improve the receptor stability. A Coomassie-stained protein gel shows that we successfully purified all mutations by IMAC to a purity of 95% (Fig 5B). Among all mutations, we found that the control A163V and M255V did not improve the purification yield or stability of FP (Fig 5C-D). In contrast, S127A exhibited a significant increase in purification yield along with a slight apparent improvement in stability, whereas A15G showed a minor apparent increase in purification yield but it did not affect receptor stability in agreement with our predictions. Overall, both S127A and A15G were found to enhance the purification outcome of FP, albeit to a different extent. We next sought to assess the impact of our mutants and deletions on FP signaling using a Rho-based BRET cellular assay that captures all cognate G proteins coupling of the receptor, including G_12/13_ [3]. The dose-response curves of PGF_2α_ and cloprostenol in HEK293 cells overexpressing FP receptors with N-terminal deletions at positions 13, 18 and 19, as well as A15G and S127A mutants, yielded EC50 and Emax values that were not statistically different from those observed with of the wild-type receptor. These findings indicate that the N-terminal deletions and the A15G and S127A mutations do not impact the receptor’s function (Fig 6).

**Fig 5.**
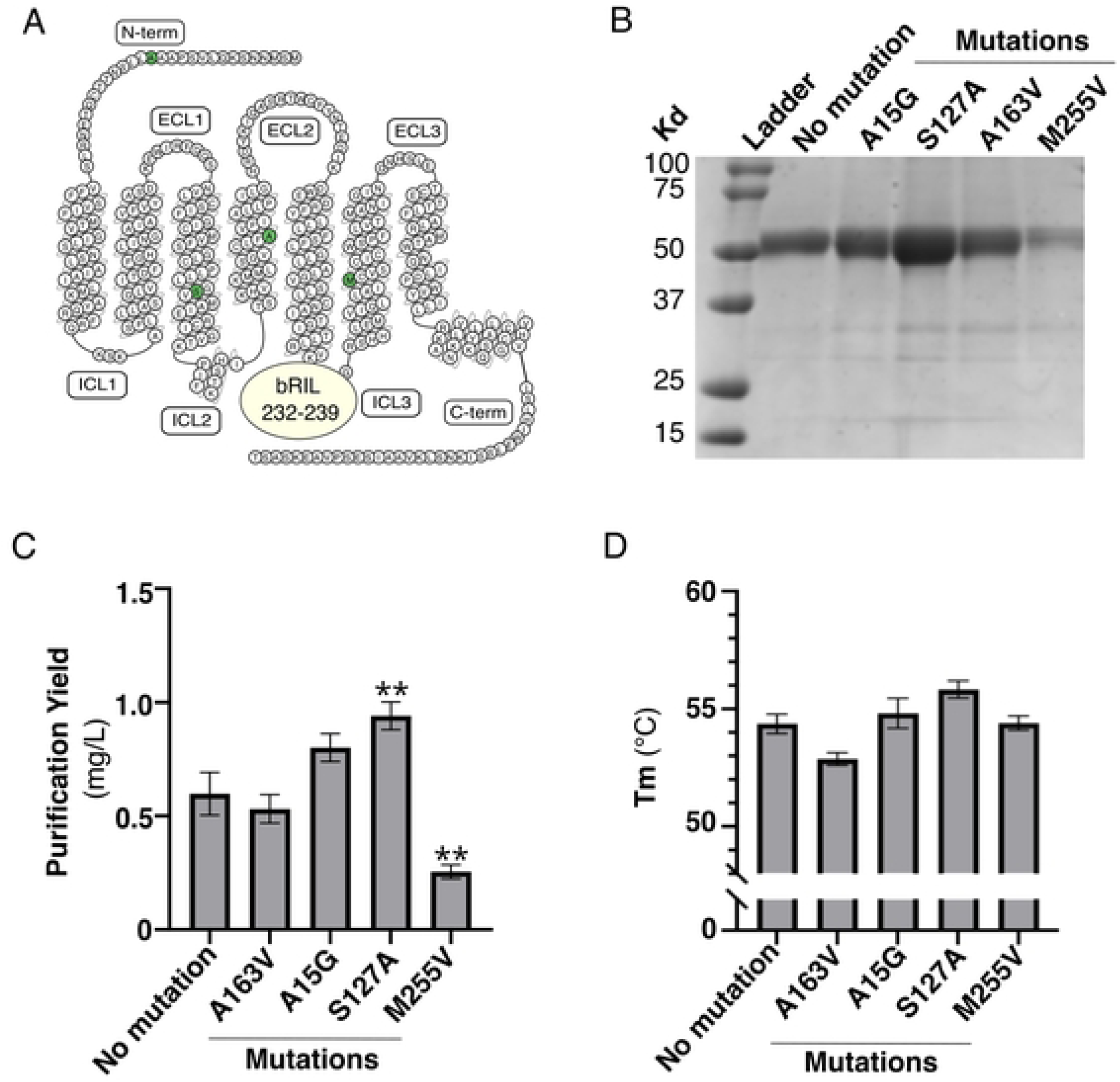
Assessment of predicted stabilizing mutations on FP. (A) Snake plot of FP. Mutations are indicated by green circles. bRIL insertion in FP ICL3 between residues 232 and 239 is shown by a circle. (B) Coomassie-stained 12% polyacrylamide gel, (C) purification yield, and (D) melting temperature (Tm) of the IMAC eluate of the indicated FP constructs. The parental construct is described in Fig 3. The data are expressed as the means ± SEM of 3 independent experiments. Statistical significance was determined by one-way ANOVA followed by a Dunnett post-hoc analysis. ** p-value < 0.01.

**Fig 6.**
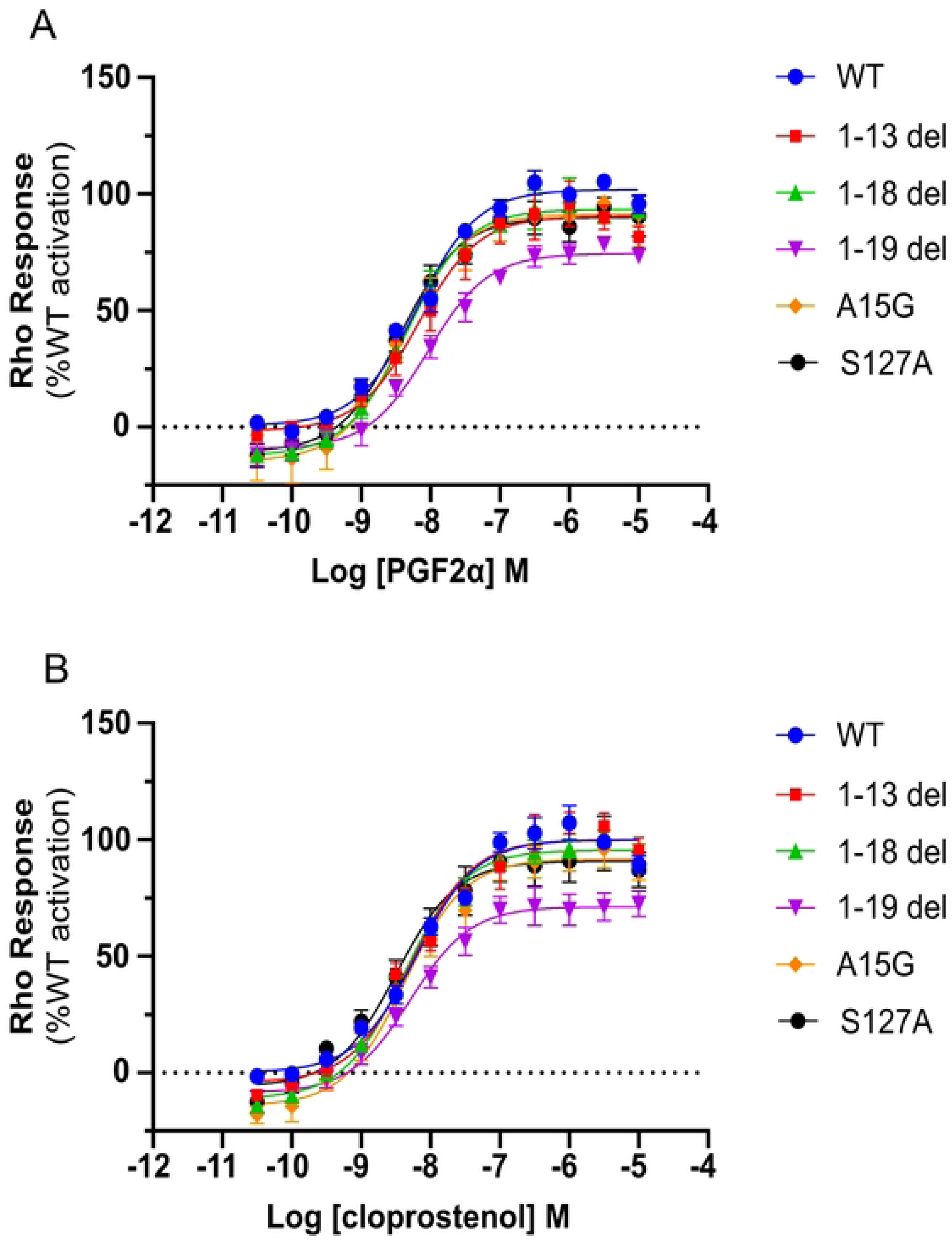
Functional impact of FP modifications. A) PGF_2α_ and B) cloprostenol-induced FP response on the Rho signaling pathway. WT and mutant FP receptors were expressed in HEK 293T cells along with the RhoA BRET biosensor to measure G protein response following PGF_2ɑ_ and cloprostenol stimulation, respectively. BRET response was normalized as a percentage of the WT maximal response and 12-point concentration-response curves were plotted using GraphPad Prism (see Methods). The data are expressed as the means ± SEM of A) 4 and B) 5 independent experiments. Statistical significance was determined by paired Student’s T test and correcting for multiple comparisons using the Holm-Šídák method. *p<0.05.

### Screening of small soluble domains and combination of most stable modifications

Our previous screening of bRIL fusion within the FP ICL3 allowed us to identify a few key insertion sites for small soluble domains that improve the purification yield and stability of FP-WT (Fig 2). To evaluate the potential benefit of the fusion of other small soluble domains previously used to stabilize GPCRs, we inserted T4 lysozyme (T4L), the thermostable glycogen synthase domain from *Pyrococcus abyssi* (PGS), rubredoxin, flavodoxin or xylanase between residues 233-239 of FP (Fig 7A). We also included the favorable mutations A15G and S127A in all FP fusions [18, 20, 29]. The new constructs were successfully expressed, and FP-fusions eluates were obtained at more than 95% purity (Fig 7B). Rubredoxin and Xylanase fusions led to an improvement in FP purification yield, with the rubredoxin fusion displaying the best purification yield at approximately 1.5 mg/L (Fig 7C). In contrast, T4L and PGS fusions decreased the observed yield, and did not improve the stability of FP (Fig 7C-D). Although the insertion of flavodoxin did not affect the purification yield of FP, both rubredoxin and flavodoxin fusions augmented the stability of FP with a Tm of 58 and 57°C, respectively. Overall, the rubredoxin fusion with FP displayed the best improvement in purification yield and stability compared to FP with a bRIL insertion.

**Fig 7.**
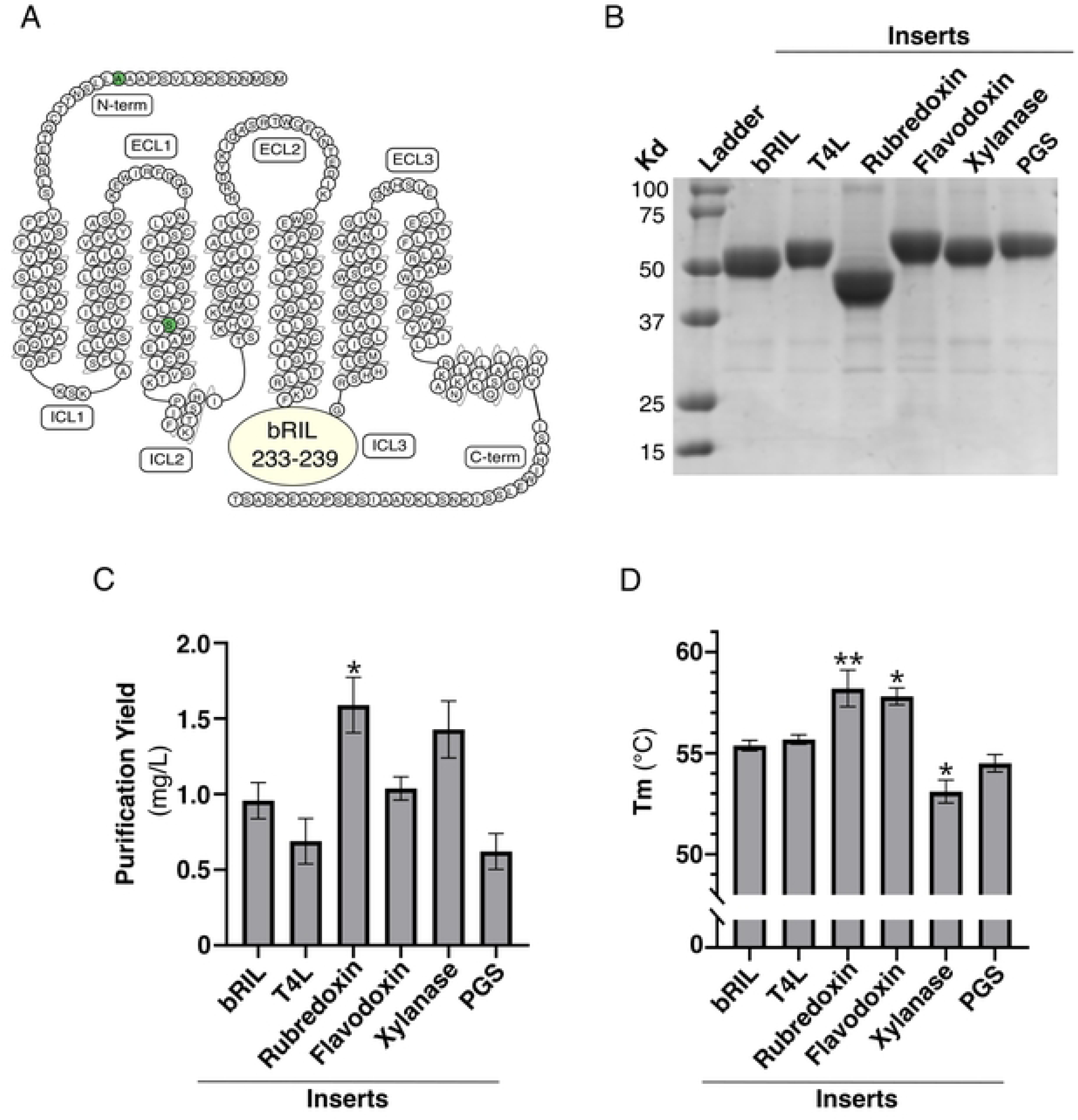
Impact of soluble domain on FP. (A) Snake plot of FP. FP fusions have the indicated small domain inserted between residues 233-239 of the ICL3, and harbor stabilizing mutations A15G and S127A (green circles). (B) Coomassie-stained 12% polyacrylamide gel, (C) purification yield and (D) melting temperature (Tm) of the IMAC eluate of the indicated FP fusions with different soluble domains. The data are expressed as the means ± SEM of 3 independent experiments. Statistical significance was determined by one-way ANOVA followed by a Dunnett post-hoc analysis. * p-value < 0.05; **; p-value < 0.01.

Finally, we took advantage of the modifications improving FP purification yield and stability and opted to combine a few of the most effective ones (Fig 8). Although we found that N-terminal and C-terminal truncations display similar purification yield and stability compared to the untruncated receptor, removal of a flexible domain of the receptor could potentially prevent aggregation at the high concentration required for crystallography and cryoEM, as well as promote the nucleation process in crystallography. Thus, we have selected two of the least disrupting truncations, deletions of the N-terminus at residues 13 and 19, to assess their impact on FP constructs’ purification yield and stability, combining optimal fusions and mutations. We present the results for three new constructs that improve the rubredoxin fusion found in Fig 7. In addition to the N-terminal truncations, these constructs display rubredoxin insertion between residues 232-239 or 233-239 of FP ICL3, and the mutation S127A (Fig 8A). We successfully purified these constructs at a purity of more than 95% (Fig 8B). The purification yield was 1.5 mg/L for all the constructs, a result similar to the rubredoxin fusion of Fig 7 (control in Fig 8). In contrast, they displayed a 2°C improvement of their Tm, suggesting that combining optimal small domain insertions, mutations, and N-terminal truncation leads to a stabilization of the constructs. Together, we show that we were able to improve the purification yield of FP-cloprostenol complex by approximately threefold and the stability of the micelles-containing receptor by 9°C, leading to a protein preparation suitable for further structural biology studies.

**Fig 8.**
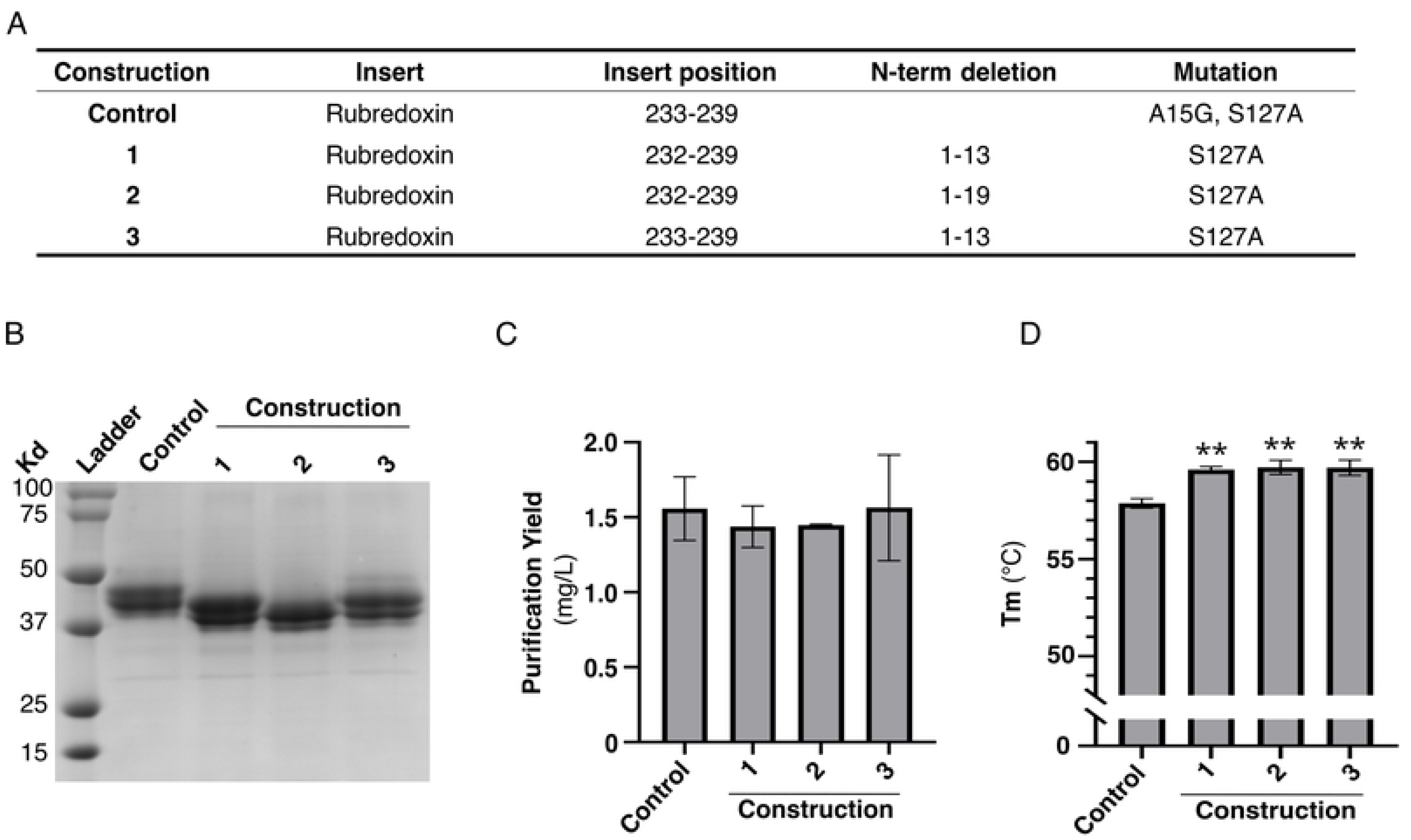
Optimized FP constructs. (A) Description of stable constructs. (B) Coomassie-stained 12% polyacrylamide gel, (C) purification yield and (D) melting temperature (Tm) of IMAC eluate of indicated FP constructs. The data are expressed as the means ± SEM of 3 independent experiments. Statistical significance is determined by one-way ANOVA followed by a Dunnett post-hoc analysis. ** p-value < 0.01.

## Discussion

GPCRs are inherently unstable *in vitro,* leading to receptor aggregation at concentrations used for structural biology. Here, we reported the optimization of the protein sequence of the FP-cloprostenol complex to obtain a stable and monodispersed purified receptor using approaches such as N- and C-terminal truncation, mutations, and insertion of small soluble domains. Truncation of the unstructured N- and C-terminus is an interesting strategy in structural biology to reduce the probability of interfering with crystal contact and to improve diffraction, preventing receptor aggregation at high concentration. The C-terminal domain of GPCRs is also important for receptor desensitization in cells. It is the target of many post-translational modifications such as phosphorylation by protein kinases that induce the recruitment of ßarrestins, which act as scaffolding effectors for cellular desensitization machinery [30]. Truncation of the C-terminus of many GPCRs may increase the receptor purification yield by blocking cell surface receptor engagement of ßarrestins, and is a strategy that has been used for GPCRs optimization for structural biology [16, 19, 31, 32]. Here, truncation of FP’s C-terminal domain did not affect receptor cell surface expression and *in vitro* stability, but rather generally slightly reduced the purification yield. This correlates with previous reports showing little or no FP coupling to ßarrestins in cells and points to distinct cellular regulation mechanisms of the receptor [3, 33, 34]. Absence of ßarrestins coupling has already been reported for EP3, another prostaglandin receptor [33, 35]. Interestingly, we could not truncate the C-terminus of FP shorter than residue 324, located immediately before the two cysteines at position 322-323, without major effects on receptor stability. Many cysteine residues at the end of helix 8 of GPCRs are palmitoylated. This post-translational modification serves as a membrane anchor for helix 8 and is involved in receptor folding and trafficking [36]. Although palmitoylation of FP has never been described before, our findings suggest the critical importance of helix 8 between residues 308-323 for FP stability *in vitro*. Interestingly, cysteine 323 points toward the membrane in the FP cryoEM structure, suggests that the C-terminus of helix 8 could be palmitoylated and that the palmitoyl-cysteines could play an important role in FP stability. Another efficient strategy for stabilizing GPCRs is the introduction of mutations, and single point mutation have shown to stabilize GPCRs for crystallization. For example, replacement of S91 by a lysine leads to a significant thermostabilization of the adenosine 2A receptor [37]. Similarly, introduction of two mutations enabled the crystallization of the serotonin 2A receptor [32]. In our study, replacement of the membrane-exposed polar serine 127 by alanine, a closely related hydrophobic amino acid, probably tightens the hydrophobic bonds of transmembrane helix 3 with the micelles leading to the increase of FP thermostability *in vitro*. Additional thermostabilizing mutations could be identified from the agonist-bound FP cryoEM structures [14, 15]. These mutations could potentially enable the study of other FP ligands with different efficacies.

Insertion of small soluble domain is one of the most important modifications present in many of the GPCR structures. Here, we used the small soluble domain bRIL to screen for the optimal ICL3 insertion site. We then used the best site to screen for other small soluble domains, as they have been designed initially with similar geometrical requirements for GPCR loop insertion [18]. However, there are different structural constraints among these domains. For example, the N-terminus of bRIL is an alpha helix that dictates its orientation within the fusion protein relative to the receptor. Thus, it is possible that the insertion site would differ slightly among the small soluble domains and re-optimization could be performed for each insert after the initial screen. Alternatively, the introduction of glycine-serine linker on each side of the insert has already been used to circumvent this challenge [17].

The receptor engineering strategies used in our study apply to the two main structural biology approaches to solve GPCR structures at high-resolution that are cryoEM and X-ray crystallography. Typical concentrations of purified GPCR needed for cryoEM and X-ray crystallography are similar and range from 2-5 and 20-30 mg/ml, respectively. Although protein concentration required for cryoEM is lower, both approaches require a similar amount of initial receptor due to the difficulty in cryoEM to obtain optimal vitrification conditions. Thus, a minimum purification yield of 0.3-0.5 mg/L of stable receptor is required for screening and collecting multiple EM grids, as well as for initial screens in X-ray crystallography [24]. Although cryoEM typically requires less receptor modification, most structures are agonist-bound due to the necessity to co-purify the receptor in a complex with the heterotrimeric G proteins and nanobodies to respect the mass limitation and circumvent the particle orientation problem. The presence of heterotrimeric G protein would likely shift the receptor equilibrium toward an activated state, displaying a structure with a higher affinity for agonists. Thus, obtaining an antagonist-bound cryoEM co-structure remains a challenge [38]. To date, X-ray crystallography has provided a greater diversity of co-structures with ligand of different efficacy due to the protein optimization effort to obtain receptor crystals [26]. Generally, the isolation of antagonist-receptor complexes within the prostaglandin receptor subfamily is notoriously challenging. The only available antagonist-bound co-structure is for the PGE_2_ type 4 receptor [21]. Similarly, FP is not stable in its apo form and in complex with an antagonist *in vitro*. Thus, we used the agonist cloprostenol as a stabilizing tool to obtain initial IMAC eluates of FP. It is possible that the impact of FP modifications would be specific for agonists as antagonist co-purification with our optimal FP constructs did not yet yield to stable IMAC eluates. Thus, solving the structure of antagonist-bound FP using cryoEM or X-ray crystallography approaches would require further receptor optimization. As such, our optimized FP constructs display a stability and purification yield well within structural biology working parameters and constitute a good initial framework for further receptor optimization.

## Acknowledgments

The authors thank all the staff at the Pharmacology Institute of Sherbrooke for their assistance.

## Supporting information

**S1 Table. Expression of the FP constructs in *Sf*9 cells.** Cell expression was monitored by flow cytometry as previously described [24]. The data are represented as the means ± SEM of 3 independent experiments.

